# Nonsense-associated alternative splicing as a putative reno-protective mechanism in *Pkhd1^cyli^/Pkhd1^cyli^* mutant mice

**DOI:** 10.1101/2021.09.22.461297

**Authors:** Chaozhe Yang, Naoe Harafuji, Maryanne C. Odinakachukwu, Ljubica Caldovic, Ravindra Boddu, Heather Gordish-Dressman, Oded Foreman, Eva M. Eicher, Lisa M. Guay-Woodford

**Author notes:** Corresponding author (LG-W).

## Abstract

Autosomal recessive polycystic kidney disease (ARPKD) is a hereditary hepato-renal fibrocystic disorder and a significant genetic cause of childhood morbidity and mortality. Mutations in the Polycystic Kidney and Hepatic Disease 1 (*PKHD1*) gene cause all typical forms of ARPKD. Several mouse strains carrying diverse genetically engineered disruptions in the orthologous *Pkhd1* gene have been generated. The current study describes a novel spontaneous mouse recessive mutation causing a cystic liver phenotype resembling the hepato-biliary disease characteristic of human ARPKD. Here we describe mapping of the cystic liver mutation to the *Pkhd1* interval on Chromosome 1 and identification of a frameshift mutation within *Pkhd1* exon 48 predicted to result in premature translation termination. Mice homozygous for the new mutation, symbollzed *Pkhd1*^*cyli*^, lack renal pathology, consistent with previously generated *Pkhd1* mouse mutants that fail to recapitulate human kidney disease. We have identified a profile of alternatively spliced *Pkhd1* renal transcripts that are distinct in normal versus mutant mice. The *Pkhd1* transcript profile in mutant kidneys is consistent with predicted outcomes of nonsense-associated alternative splicing (NAS) and nonsense mediated decay (NMD). Overall levels of *Pkhd1* transcripts in mutant mouse kidneys were reduced compared to kidneys of normal mice, and *Pkhd1* encoded protein in mutant kidneys was undetectable by immunoblotting. We suggest that in *Pkhd1*^*cyli*^*/Pkhd1*^*cyli*^ *(cyli)* mice, mutation-promoted *Pkhd1* alternative splicing in the kidney yields transcripts encoding low-abundance protein isoforms lacking exon 48 encoded amino acid sequences that are sufficiently functional so as to attenuate expression of a renal cystic disease phenotype.

## Introduction

Autosomal recessive polycystic kidney disease (ARPKD; MIM263200) is a hereditary hepato-renal fibrocystic disorder with an estimated incidence of 1 in 26,500 live births [1]. ARPKD is characterized by the formation of renal cysts affecting the collecting ducts, causing progressive renal insufficiency and ultimately end stage kidney disease in most patients [2, 3]. The disease also affects the liver with biliary plate malformations leading to portal hypertension and hepatic fibrosis [2, 3]. Virtually all cases of typical ARPKD are caused by mutations within the polycystic kidney and hepatic disease 1 (*PKHD1*) gene, located on chromosome 6p21.1 [4-6]. The full-length *PKHD1* transcript is composed of 67 exons with the longest open reading frame (ORF) encoding a 4,074 amino acid protein called fibrocystin or fibrocystin/polyductin complex (FPC) [4, 5]. Despite the identification of *PKHD1* as the genetic determinant of ARPKD almost two decades ago, the function of FPC remains undefined.

Several orthologous mouse models of ARPKD have thus far been described (**Table 1**), primarily generated through random mutagenesis or targeted genetic engineering of the *Pkhd1* gene [7-14]. Most mutant *Pkhd1* mice exhibit a liver phenotype resembling human disease, but kidney cystic disease is either absent or very mild and slowly progressive [3]. The mouse *Pkhd1* locus, located on Chromosome 1qA3-4, consists of 67 non-overlapping exons encoding a protein of 4,059 amino acids [15]. Human and mouse FPC share 87% overall identity across domains encompassing a predicted N-terminal signal peptide, multiple immunoglobulin-like plexin domains, multiple parallel β-helix 1 repeats and a single transmembrane domain. In contrast, the orthologous proteins share short C-terminal cytoplasmic domains that are only 40% identical [15, 16].

**Table 1.** ARPKD orthologous mouse model phenotypes.

| Strain | Background | Kidney | Liver | Pancreas |
| --- | --- | --- | --- | --- |
| *Pkhd1<sup>lacZ</sup>* | 129S; B6 | PT dilatation | DPM | Cystic |
| *Pkhd1<sup>LSL(-),Pk(+)</sup>* | BALB/cJ, C57BL/6 | PT dilatation | DPM | Cystic |
| *Pkhd1<sup>del2</sup>* | BALB/cJ, C57BL/6 | PT dilatation | DPM | Cystic |
| *Pkhd1<sup>del3-4</sup>* | 129S; B6 | TAL/CD dilatation | DPM | Cystic |
| *Pkhd1<sup>del4</sup>* | 129S; B6 | None | DPM | Cystic |
| *Pkhd1<sup>e15GFPdel16</sup>* | C57BL/6 | PT/MCD dilatation | DPM | None |
| *Pkhd1<sup>del40</sup>* | C57BL/6 | None | DPM | ---- |
| *Pkhd1<sup>cyli</sup>* | C57BL/6 | None | DPM | None |
| *Pkhd1<sup>del67</sup>* | C57BL/6 | None | None | None |

PT, Proximal tubule; TAL, thick ascending limb of Henle; MCD, medullary collecting duct; CD, collecting duct; DPM, ductal plate malformation.
Mutation types and references: *Pkhd1<sup>lacZ</sup>*, exons 1-3 replaced by nLacZ/Neo insertion [12]; *Pkhd1<sup>LSL(-)</sup>*, LoxPSTOPLoxP (LSL) insertion in exon 2 & *Pkhd1<sup>Pk(+)</sup>*, Cre-mediated recombination of LSL [17]; *Pkhd1<sup>del2</sup>* exon 2 deletion [8]; *Pkhd1<sup>del3-4</sup>* deletion of exons 3 and 4 [9]; *Pkhd1<sup>del4</sup>* exon 4 deletion[10], *Pkhd1<sup>e15GFPdel16</sup>* disruption of exon 15 and deletion of exon 16 [11], *Pkhd1<sup>del40</sup>* disruption of exon 40 leading to exon 40 “skipping” [7], *Pkhd1<sup>del67</sup>* exon 67 deletion [14].

Numerous alternative *Pkhd1* transcripts have been reported [15-17] while the inventory of alternatively spliced human *PKHD1* transcripts appears to be less complex [18]. More than 20 alternative *Pkhd1* transcripts were identified in wild-type (normal) mice [15, 16]. Several *Pkhd1* intronic and exonic splicing enhancers essential for proper *Pkhd1* splicing *in vitro* have been described [16]. Database analysis of *PKHD1* missense mutations associated with ARPKD (http://www.humgen.rwth-aachen.de) has identified sequence variants predicted to disrupt normal splicing, leading to premature protein termination [16].

Here we report discovery of the *Pkhd1*^*cyli*^ mutation (hereafter, symbolized *cyli* for ease of presentation). We describe the phenotype of *cyli*/*cyli* mice, the mapping and identification of the disease gene, as well as comparative studies of *Pkhd1* transcript profiles and abundance in normal and *cyli/cyli* mice. The phenotype in *cyli/cyli* mice is consistent with what is observed in most other orthologous mouse models of human ARPKD; a largely liver-restricted cystic disease lacking renal involvement. The *cyli* mutation is an indel in exon 48 that results in a frameshift leading to premature protein termination. Kidney *Pkhd1* transcript profiles differed both qualitatively and quantitatively between normal and *cyli/cyli* mutant mice. Taken together, our observations suggest that the *cyli* mutation results in the activation of both nonsense-associated alternative splicing (NAS) [20-22] and nonsense-mediated decay (NMD) [22-27] mechanisms in mutant kidneys. We propose that in the *cyli/cyli* mouse, the absence of renal cystic disease is due to a combination of nonsense-associated alternative splicing (NAS) [20-22] that generates *Pkhd1* mRNAs lacking mutated exon 48, thereby avoiding premature protein termination, and NMD eliminating the majority of normally-spliced exon 48-containing transcripts. We suggest that the resulting *Pkhd1* transcript pool directs translation of FPC isoforms of low abundance but sufficient function to attenuate expression of a renal disease phenotype.

## Materials and Methods

### Mice

All protocols were approved by the Animal Care and Use Committees at The Jackson Laboratory, University of Alabama at Birmingham (UAB) and Children’s National Research Institute. The study was conducted in accordance with the recommendations in the Guide for the Care and Use of Laboratory Animals of the National Institutes of Health. The Jackson Laboratory, UAB, and the Research Animal Facility at Children’s National Medical Center are fully accredited by the AAALAC.

The D.B/11Ei congenic mouse strain was generated by introgression of a segment of distal Chromosome 4 from C57BL/6J (B6) onto the DBA/2J background. The first affected mouse noted was a 5 month-old D.B/11Ei female breeder, generation N11F13. This female had successfully raised two litters, was pregnant with a third litter, and appeared sick. Further investigation revealed a hardened, distended abdomen containing an enlarged liver with yellow, fluid-filled cysts. Liver disease was not evident in the male breeder. Offspring from this pair and closely related mice were monitored for signs of liver disease. These offspring were used to establish the D.B/11Ei strain carrying the defective gene and the results reported here derive from the original breeding pair.

The mice used in this study were first transferred to UAB and subsequently to Children’s National Research Institute. Because affected mice survive into adulthood and are capable of reproducing, the *cyli* mutation is maintained in the D.B/11Ei homozygous *cyli* breeding colony.

### Locus mapping, gene identification and mutation sequencing

A standard backcross mating scheme was used to identify the chromosomal location of the *cyli* gene [28-30]. F_1_ females, produced by mating B6 to D.B/11Ei-*cyli* mice, were backcrossed to D.B/11Ei-*cyli* males. The backcross offspring (n = 221) were evaluated at 5 months of age for the presence of cystic liver disease. An initial genetic variant mapping approach and subsequent fine mapping studies were performed using MIT microsatellite markers [31]. The introgressed B6 segment on Chromosome 4 was excluded as a candidate disease interval. A disease associated interval identified on Chromosome 1 contained the *Pkhd1* locus, which was analyzed by DNA sequencing. *Pkhd1* exons and flanking intronic sequences were PCR amplified and the amplicons bi-directionally sequenced using primer sets (Supplementary Table 1) designed based upon the published sequence [15].

### Mouse genotyping

DNA for genotyping was isolated from biopsied tail tissue. Tissue was lysed at 55°C in Cell Lysis Solution (Qiagen) containing Proteinase K (Qiagen), followed by protein precipitation with the Protein Precipitation Solution (Qiagen) for 10 min at -20 °C. The sample was then centrifuged at 16,000 x g for 10 min at 4 °C. Genomic DNA was precipitated from the supernatant by addition of ethanol, pelleted by centrifugation at 16,000 x g for 5 min at 4 °C, air dried and resuspended in water. PCR-based genotyping was performed using primers 5’-TGG CTA TAC TGT GAA GAC CAG GCA-3’ (forward) and 5’-AAG CTT GGG CCT ATC TGA ATG GCA-3’ (reverse) and the following conditions: 15 min at 95 °C initial denaturation, followed by 35 cycles of 45 sec denaturation at 94 °C, 45 sec annealing at 52 °C and 1 min extension at 72 °C; with a 10 min final extension at 72 °C. PCR products were digested with *Bsa*I and the products resolved by agarose gel electrophoresis. Bands of 126 bp and 359 bp were diagnostic of normal genotype. A 484 bp band identified the *cyli* mutant allele.

### Tissue histology and morphometric analysis

Kidneys and livers harvested from 1, 2, 4 and 6 month old male and female normal and *cyli* mice were paraffin embedded, sectioned, deparaffinized, rehydrated and stained with hematoxylin and eosin according to standard protocols [32-35]. Stained tissue sections were examined by light microscopy using an Olympus CX41 microscope equipped with a Leica DX320 color camera using Leica software. Histomorphometry was performed on blinded experimental specimens by a veterinary pathologist according to previously described protocols [36]. Images were collected using a Nikon E600 microscope equipped with a SPOT Insight digital camera (Diagnostic Instruments) and analyzed using Image-Pro Plus v6.2 image analysis software (Media Cybernetics Inc). Cyst and tissue areas were quantified by converting images to gray-scale and thresholding them to produce a black image on a white background. Cysts were represented as white objects within the image. Cystic and total (including cysts) areas were determined automatically using the count/size and macro functions of Image-Pro Plus. The results were expressed as % of cyst area relative to total area.

To investigate the course of disease in D.B/11Ei-*cyli* mice, the mice were weaned at 3 weeks of age and assigned to a specific age group. To investigate liver disease progression in females as a function of litter parity, 3 sib mated pairs were assigned to be investigated after the birth of their first litter, and 3 sib-mated pairs were assigned to be investigated after the birth of their second litter.

### Reverse transcription (RT)-PCR and quantitative (q)RT-PCR

Total RNA samples from kidneys and livers harvested from 7-week-old normal and *cyli* mice were prepared using RNeasy Mini kit (Qiagen, # 74104), treated with RQ1 RNase-Free DNase (Promega, # M6101) and then re-purified using the RNeasy Mini kit. For RT-PCR, RNA samples were reverse transcribed using SuperScript III First-Strand Synthesis SuperMix (Thermo Fisher Scientific, # 18080400) and oligo dT primers. RT-PCR to compare *Pkhd1* transcript “profiles” between normal and *cyli* kidney and liver tissue was performed using primers specific for *Pkhd1* exons 1 (forward: 5’-CAT TTG AGG CAC AAG GCT GAC ACA-3’) and 67 (reverse: 5’-CTG AGG TCT GGG CGT AAC AG-3’) sequences. Relative *Pkhd1* transcript abundance in normal vs. *cyli* kidneys was determined by quantitative real-time PCR performed on a QuantStudio 7 Flex Real-Time PCR System (Thermo Fisher Scientific) using the default program. The PCR was performed on cDNA templates using Power SYBR Green PCR Master Mix (Thermo Fisher Scientific, # 4368706) and primers specific for sequences in *Pkhd1* exons 48-49 (forward: 5’-TGG CTA TAC TGT GAA GAC CA-3’; reverse: 5’-GAT CCA AGA GCA GAG CCA TC-3’), *Pkhd1* exons 61-62 (forward: 5’-TCA CTC TTG AGA TGC CTG GC-3’; reverse: 5’-AGG TTC CCA GTT ATT AAA CTA C-3’) and *Pkhd1* exons 66-67 (forward: 5’-CCA GAA GAC ATA TCT GAA TCC CAG GC-3’; reverse: 5’-AGC AAG AGA TCC TGG AAC ACA GGT-3’). *Beta-actin* was used for normalization (forward: 5’-GGA GGG GCC GGA CTC ATC GTA CTC-3’; reverse: 5’-CCG CAT CCT CTT CCT CCC TGG AGA A-3’) [16]. Results were analyzed using QuantStudio Real-Time PCR Software and the ΔΔCt method [37].

### Immunoblotting

Kidneys were collected from 14-day old normal and *cyli* mice and immediately snap frozen in liquid nitrogen. Kidneys were homogenized on ice for 20 sec in 1 ml ice-cold RIPA buffer (Sigma-Aldrich # R0278) containing proteinase inhibitors (Protease Inhibitor Cocktail Mini-Tablet EDTA-free, Bimake # B14012). Homogenates were centrifuged for 10 min at 15,000 × g at 4 °C. BCA protein assays (Thermo Scientific # 23227) were performed on supernatants. Twenty µg of total protein was mixed with NuPAGE LDS sample buffer (Life Technologies, # NP0007) containing sample reducing agent (Life Technologies, # NP0009). Samples were heated at 100 °C for 10 min prior to electrophoresis through a Novex NuPAGE 4–12 % Bis-Tris gel (Life Technologies, # NP0335BOX) in MES SDS running buffer (Life Technologies, # NP0002) for 30 min at 200 V. Proteins were transferred to a polyvinylidene fluoride membrane using a Bio-Rad Trans-Blot Turbo Transfer System. The membrane was incubated with rat anti-mouse FPC monoclonal antibody [14] in 1× PBS plus 0.1 % Tween-20 (PBST) with 5 % bovine serum albumin overnight at 4 °C. The membrane was washed 3 times with 1× PBST, 10 min per wash, then incubated with goat anti-rat secondary antibody (Thermo Fisher Scientific # 31475 1:5000 dilution in 1× PBST with 5% non-fat dry milk) for 1 hour at room temperature, followed by 3 washes with 1× PBST. Immunoreactive bands were detected using SuperSignal West Dura chemiluminescent substrate (Thermo Fisher Scientific, # 34076) and imaged using a Bio-Rad ChemiDoc MP Imaging System.

## Results

### Discovery of the *Pkhd1*^*cyli/cyli*^ mutant mouse

Investigation of an apparently sick D.B/11 female mouse revealed an enlarged liver containing multiple fluid-filled cysts. Subsequently shown to be a stably transmitted recessive trait, the spontaneous mutation was designated cystic liver (*cyli*). Homozygous *cyli* mutant mice developed cystic liver disease by two months of age. Histopathological analysis (**Fig. 1A**) demonstrated biliary dysgenesis characterized by ductal plate malformation phenocopying the liver lesion characteristic of human ARPKD, with portal tracts of affected livers exhibiting multiple irregularly shaped and variably dilated bile ducts generally lined with a hyperplastic epithelium. Histopathologic evidence of renal cystic disease was not observed in *cyli* homozygous mice examined at one, two, four and six months of age (data not shown).

**Figure 1.**
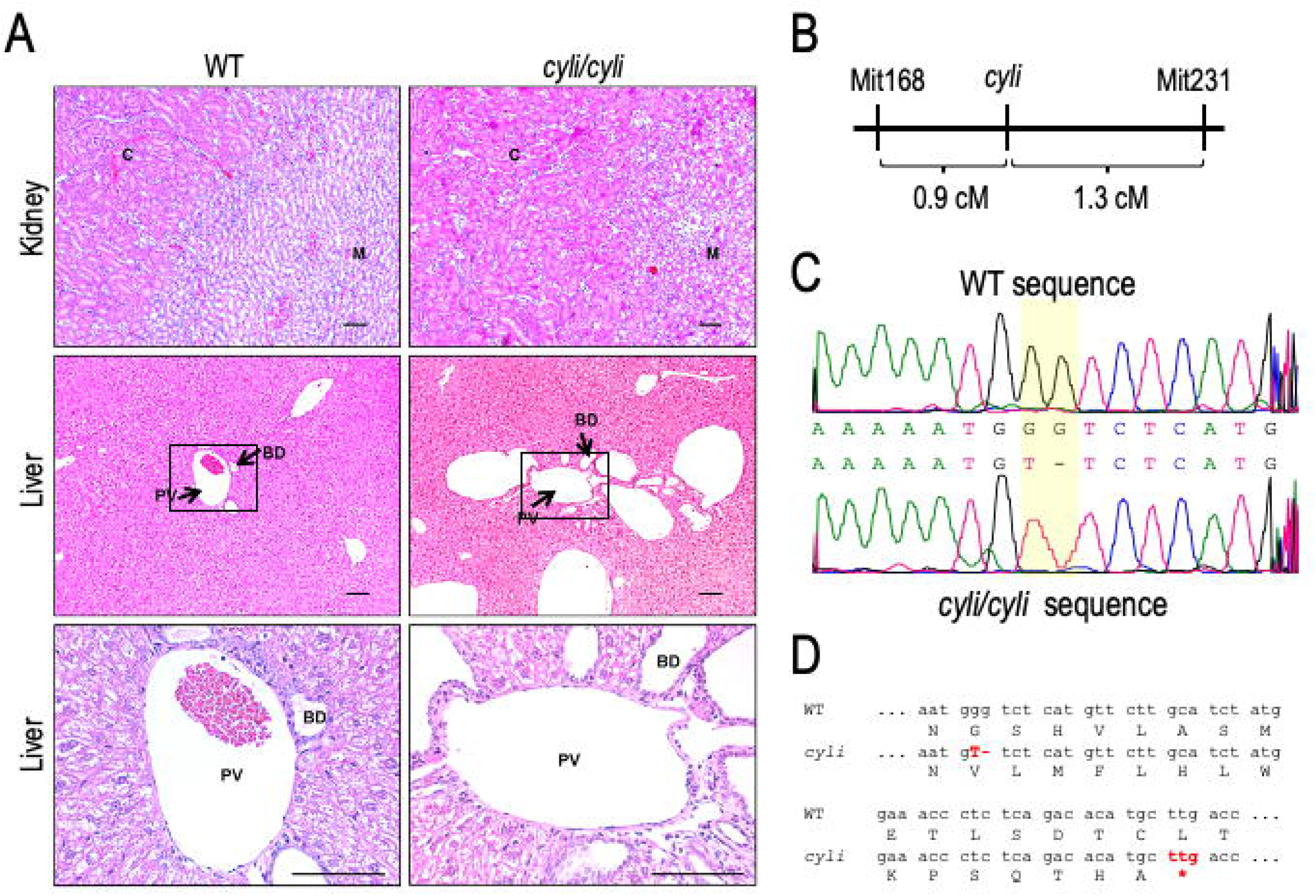
Characterization of the *cyli/cyli* mouse model. **(A)** Kidney (top) and liver (middle and bottom) tissue sections stained with hematoxylin and eosin from 2-month-old normal and *cyli/cyli* mice. Bottom panels are higher magnification views of boxed areas in middle panels (BD, Bile duct; C, cortex; M, Medulla; PV, Portal vein). Scale bar = 100 µm. **(B)** Schematic illustrating the position of the *cyli* mutation (affecting the *Pkhd1* locus) on mouse Chromosome 1 between the genetic markers *D1Mit168* and *D1Mit231* (genetic distances in centiMorgan (cM) units). **(C)** Sequence comparison of *Pkhd1* from normal and *cyli mice* (c.7588_7589delGGinsT, p. L2545fs, where nucleotide A of the translation initiation codon ATG in NM_153179.3 is +1 in *Pkhd1* exon 48). **(D)** Comparison of normal and *cyli* reading frames. Mutation leads to premature protein termination in the *cyli/cyli* mouse.

### Gene identification and mutation analysis

The disease locus was positioned on Chromosome 1 between markers *D1Mit168* and *D1Mit231* (*D1Mit168* - 0.9 cM – *cyli* - 1.3 cM – *D1Mit231*), in an interval containing the *Pkhd1* gene. (**Fig. 1B**). Sequence analysis of the *Pkhd1* gene in *cyli/cyli* mice identified a deletion/insertion mutation in *Pkhd1* exon 48; c.7588_7589delGGinsT, p. L2545fs (where nucleotide A of the translation initiation codon ATG in NM_153179.3 is +1) (**Fig. 1C**) leading to a frameshift and premature stop codon within exon 48, 44 bp downstream of the T insertion (**Fig. 1D**).

### Progressive liver disease linked to gender, age and parity

Morphometric analysis of liver sections from female and male *cyli/cyli* mice at multiple time points revealed an age-associated increase in phenotypic severity, indicating progressive liver disease. Younger *cyli/cyli* mice (less than 2-months-old) displayed a pre-cystic liver phenotype characterized by dilated bile ducts radiating from the portal region into the parenchyma (**Fig. 1 and Table 2**). Older mice displayed coalescing cysts that progressively replaced increasing areas of the normal parenchyma. Beyond 4 months of age, female mice tended to exhibit more extensive cystic lesions than male mice of comparable age (**Table 2**). Severity of the liver phenotype in females also was correlated with increased parity (**Fig. 2 and Table 2**). The current study did not extend phenotypic examination of mice beyond 6 months of age.

**Table 2.**
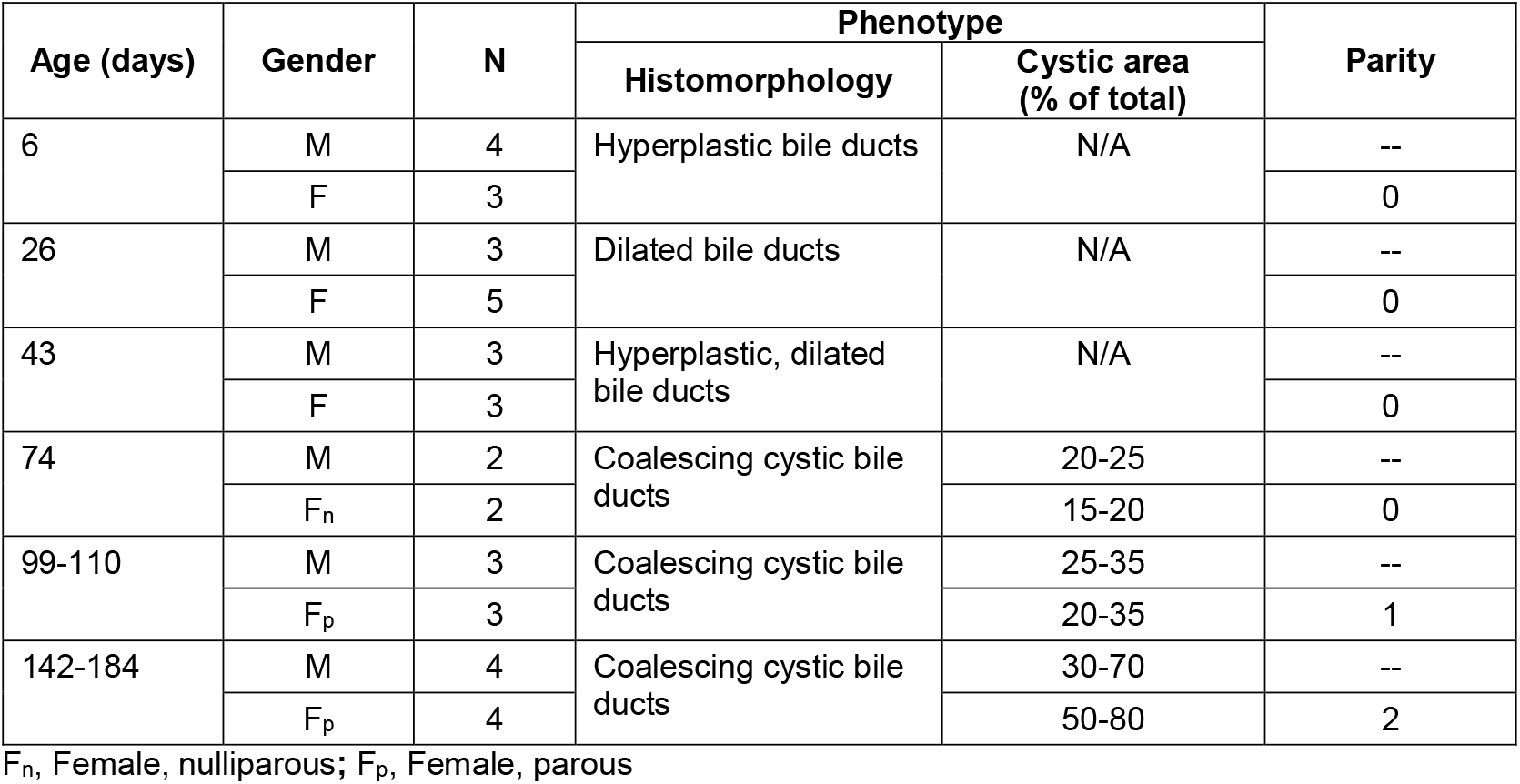
Progressive cystic liver disease in *Pkhd1*^*cyli/cyli*^ mice.

**Figure 2.**
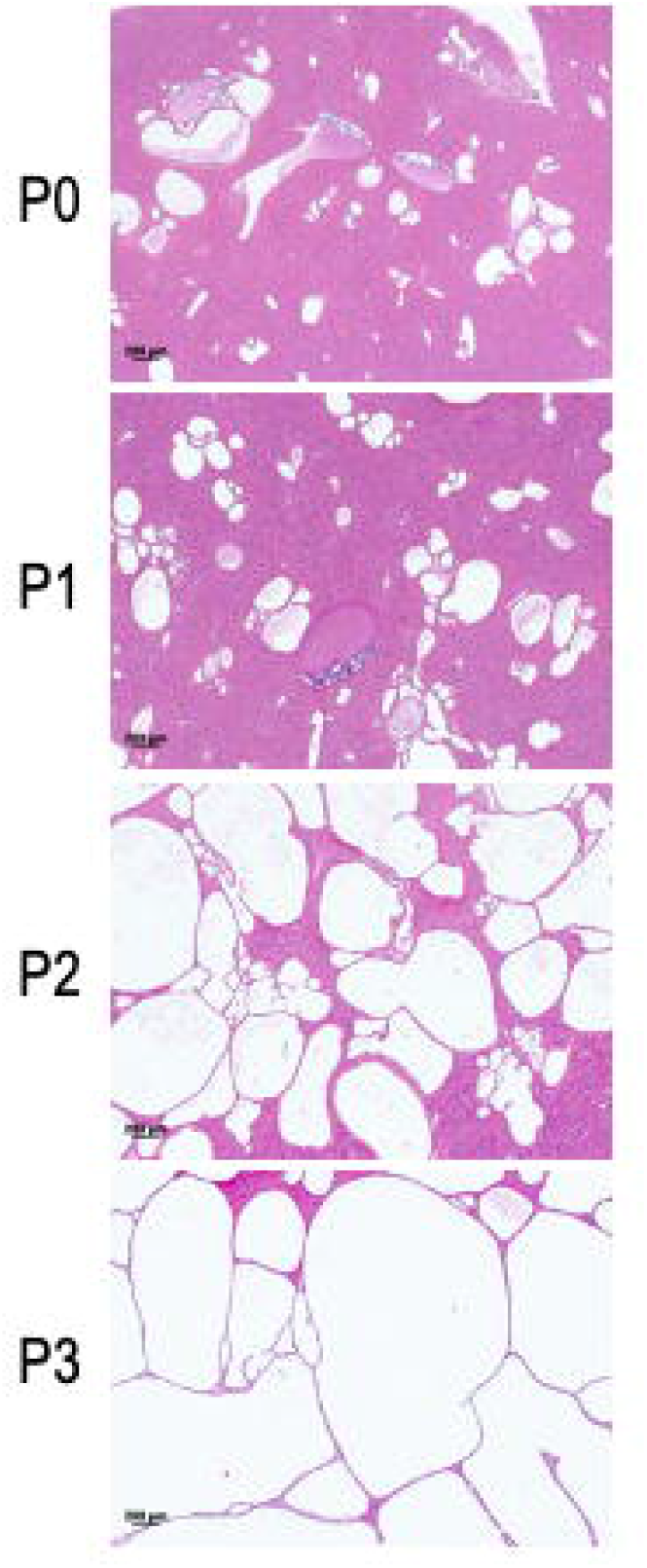
Progressive cystic liver phenotype in *cyli/cyli* mutants. Hematoxylin and eosin stained liver sections from representative *cyli/cyli* female mice, where P#0, P#1, P#2 and P#3 denote numbers of litters produced by mutant females. Scale bar = 200 µm.

### Differential *Pkhd1* transcript profiles in normal vs. *cyli/cyli* mice

There are relatively few alternatively spliced human *PKHD1* transcripts. In contrast, non-mutant mouse *Pkhd1* is subject to extensive alternative splicing in the kidney but not in the liver, and longer transcripts typically include exon 48 (location of the *cyli* mutation) [16].

Therefore, we compared the profiles of *Pkhd1* transcripts amplified from *cyli/cyli* and normal kidneys and livers (**Fig. 3A**) using RT-PCR and primers specific for *Pkhd1* exons 1 and 67. Consistent with previously published findings [16], we identified four major amplification products of 12, 6.5, 4.5 and 2.5 kbp (plus some additional minor bands) representing the full-length and alternatively spliced *Pkhd1* mRNAs from normal kidneys (**Fig. 3A, lane 3**) [16]. In normal liver, again in accordance with previous observations [16], we observed a 12 kbp amplification product (**Fig. 3A, lane 4**) while cDNA derived from the *cyli/cyli* liver yielded a 12 kbp amplicon as well as a 4.5 kbp band (**Figure 3A, lane 2**). The presence of this 4.5 kbp band is consistent with previous observations of a *Pkhd1* derived amplicon of this size from both normal liver and kidney [16]. Our RT-PCR analysis of normal and *cyli/cyli* liver samples inconsistently generated this amplicon, possibly reflecting very low transcript abundance.

**Figure 3.**
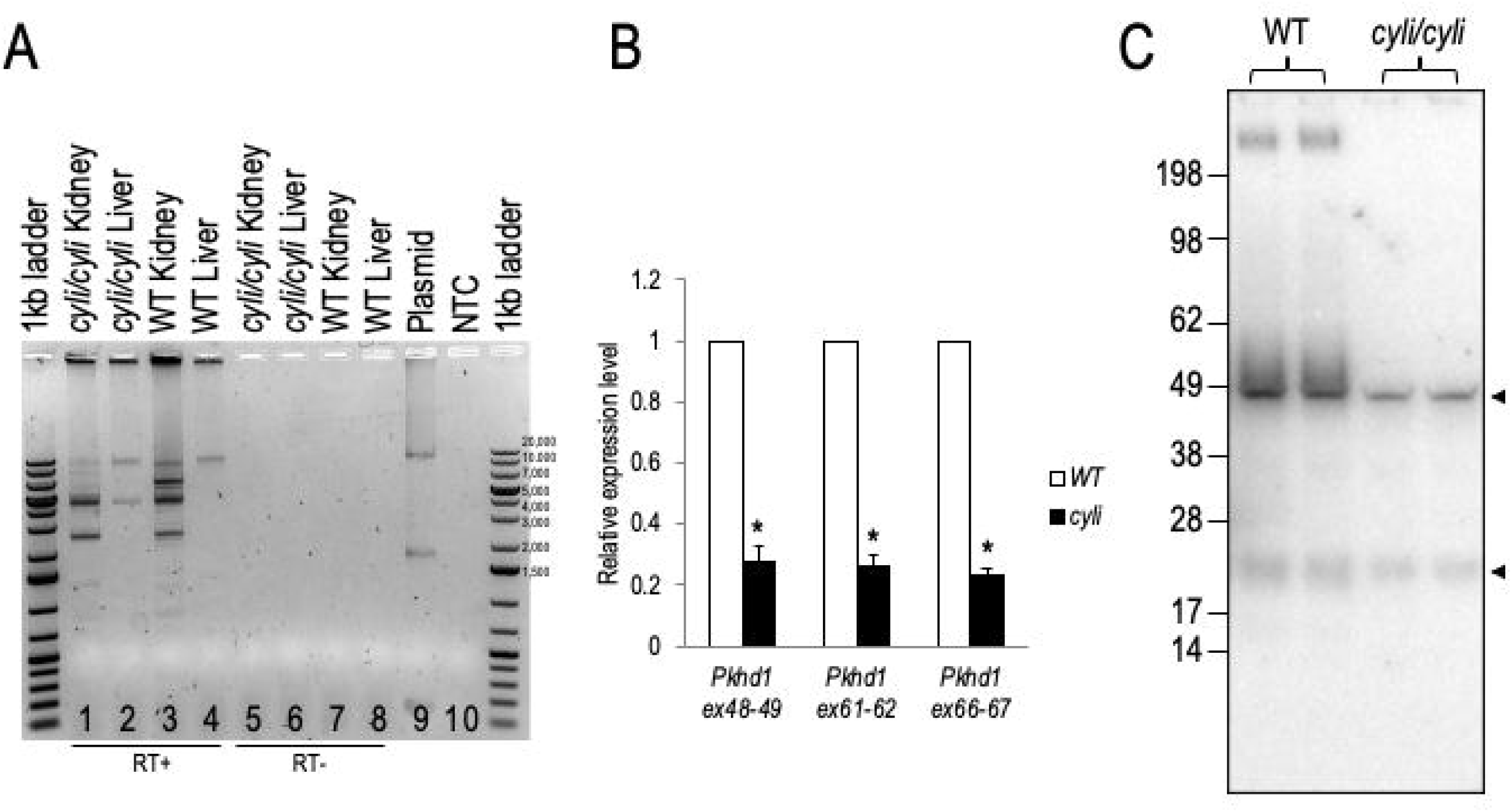
*Pkhd1* expression in normal and *cyli*/*cyli* mice. **(A)** *Pkhd1* transcript profiles in liver and kidneys of normal and *cyli*/*cyli* mice represented by PCR products (amplicons) generated from oligo-dT primed template cDNA using primers specific for *Pkhd1* exons 1 and 67. Smaller bands in lane 9 (Plasmid) represent nonspecific or plasmid recombination-derived amplification products. **(B)** Relative expression of *Pkhd1* mRNA containing exons 48-49, 61-62 and 66-67 in the kidneys of normal and *cyli*/*cyli* mice. Data were normalized to *beta actin* mRNA; expression of *Pkhd1* mRNA in normal mouse kidneys was set as 1.00. Data are expressed as mean ± S.E; n=5 per group. Statistical analysis was performed using a non-parametric Wilcoxon sign rank test. *P< 0.05 *vs*. normal. **(C)** Western blot detection of normal and *cyli*/*cyli* mouse kidney FPC protein detected using a monoclonal primary antibody specific for *Pkhd1* exon 67 encoded amino acid sequences. Only full-length FPC was observed in normal kidney protein extracts, and no FPC was detected in extracts of *cyli*/*cyli* kidneys. Results of duplicate experiments are shown. Low molecular weight bands (arrowheads) correspond to IgH and IgL chains detected by anti-mouse secondary antibody reagent.

Sequence analysis of the kidney amplicons indicated that while most transcripts represented in the 12 and 6.5 kbp bands included exon 48, this exon was excluded in transcripts represented in bands of 4.5 and 2.5 kbp. As noted, the *cyli/cyli* kidney yielded only two predominant amplification products of 4.5 and 2.5 kbp, along with a relatively faint 12 kbp and additional minor bands (**Fig. 3A, lane 1**). We used qRT-PCR and primer-pairs specific for junctions spanning exons 48-49, 61-62 and 66-67 and examined whether the differential amplicon profiles reflected exclusion of exon 48 from the population of *cyli/cyli* derived mRNAs. We observed significantly lower levels of these targeted amplicons in *cyli/cyli vs*. normal kidneys (**Fig. 3B**), consistent with NMD activity. In addition, detection of *Pkhd1*^*cyli*^ amplicons using an exon 48 specific primer indicates that NAS did not result in a complete absence of mutant exon 48 containing transcripts. Therefore, we propose that a combination of NAS and NMD-mediated mechanisms may differentially enhance the proportion of *Pkhd1* transcripts lacking exon 48 in *cyli/cyli* kidneys as compared to normal kidneys.

We also investigated expression of the *Pkhd1* encoded protein, FPC, in normal and *cyli/cyli* kidneys, using an antibody generated against an exon 67 encoded epitope (rat monoclonal antibody PD1E1, kindly provided by the Baltimore PKD Center) (**Fig. 3C**). This antibody detected full-length FPC in protein extracts only from normal kidneys. In contrast, the full-length FPC was not detected in extracts prepared from *cyli/cyli* kidneys, as expected given the nature of the *cyli* mutation. Possible lower molecular weight FPC isoforms also were not observed, a result consistent with our qRT-PCR findings of significantly reduced levels of *Pkhd1* mRNA in *cyli/cyli* kidneys, resulting in FPC abundance below the threshold level of immunoreagent detection.

## Discussion

The mouse *cyli* mutation arose spontaneously due to a *de novo* deletion/insertion (c.7588_7589delGGinsT) in exon 48 of the *Pkhd1* gene, which is predicted to be a frame shift mutation leading to premature protein termination. Similar to other *Pkhd1* gene-targeted mouse models, the *cyli/cyli* mutants express a hepato-biliary phenotype that progresses with age and in females, disease severity is accelerated by increasing parity. In the inbred D.B/11Ei congenic line, the *cyli* mutation is not associated with a renal cystic phenotype. That said, a mild renal cystic phenotype can be observed when other gene-targeted *Pkhd1* mutations are expressed on mixed genetic backgrounds or when mutant mice are aged for more than 12 months. For example, homozygotes carrying the *Pkhd1*^*C642\**^ truncating mutation do not have imaging evidence for either kidney or liver disease in the first 6-months of life. However, by ∼1.5 years of age, female *Pkhd1*^*C642\**^ heterozygotes, as well as homozygotes, develop radiographic changes resembling medullary sponge kidney. Interestingly, histopathological analysis demonstrates that the renal tubular ectasia in these mutant kidneys involves the proximal tubule rather than the collecting duct, a pattern that recapitulates the earliest ARPKD-associated renal cystic lesion in human fetuses [38].

With all the mouse models described to date, the most striking feature is the minimal renal phenotype associated with homozygous *Pkhd1* frameshift and truncating mutations. In comparison, similar human *PKHD1* mutations typically cause severe renal cystic disease that is expressed in fetuses and infants [3]. In the specific example of exon 48, the *cyli* frameshift mutation is not associated with renal cystic disease in 6-month old mice, whereas human patients with frameshift mutations involving *PKHD1* exon 48 have the classic ARPKD phenotype with renal cystic disease expressed in infancy (http://www.humgen.rwth-aachen.de) [39].

We suggest that nonsense-associated alternative splicing (NAS) may in part explain the species-specific absence of renal cystogenesis in the *cyli* model, and perhaps other engineered *Pkhd1* mutations. The mouse *Pkhd1* gene, unlike its human orthologue, is transcriptionally complex in the kidney with a number of alternatively spliced isoforms. Whereas in the liver, *Pkhd1* has minimal transcriptional complexity and the full length *Pkhd1* transcript predominates [4, 16]. In *cyli/cyli* mutant kidneys, we detected only the lower molecular weight 2.5 and 4.5 kb amplicons, consisting largely of putative transcripts lacking exon 48. Sequence analyses in a previous report [16] and data from this study demonstrate that these lower molecular weight amplicons contain isoforms that involve exon-skipping events, but maintain the FPC open reading frame (ORF). For example, the 2.5 kb amplicon contains a putative isoform with splicing from exon 6 to 61; similarly, among the transcripts in the 4.5 kb amplicon, alternative splicing events involve exon 4 to 49 and exon 6 to either 51, 52 or 53 (**Fig. 4**). Therefore, a diverse set of alternative *Pkhd1* splice forms, generated by NAS, could encode multiple novel isoforms of FPC with sufficient residual function to impede renal cystogenesis in mutant mice. In contrast, the mutant liver with limited alternative splicing would not have similar functional redundancy, resulting in development of the hepato-biliary lesion. In addition, amplicons from the *cyli/cyli* kidney did not totally exclude exon 48, suggesting the possibility that cryptic splice site(s) could be activated within the exon, resulting in a lower expression of transcripts containing exon 48, but maintaining the ORF and thus evading NMD mechanisms.

**Figure 4.**
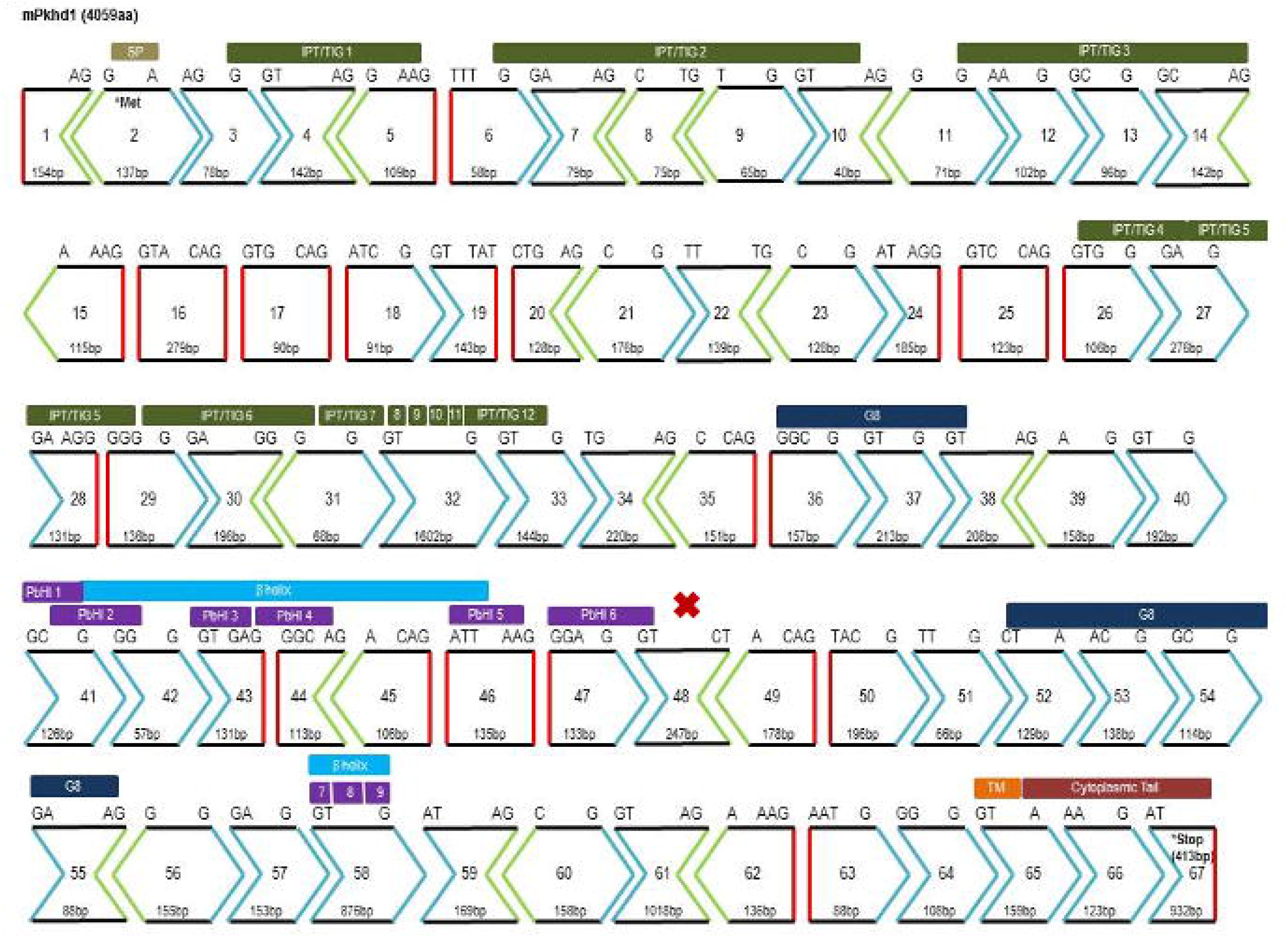
*Pkhd1* transcript structure. Schematic shows the exons in the longest ORF encoding major functional protein coding domains. The ORF is preserved when splicing occurs between exons with similar configurations (like-to-like colored/shaped exon boundaries), e.g. exon 6 to 7; exon 6 to 48, which could generate both the longest ORF transcript and alternatively spliced transcripts. The *cyli* mutation and resulting downstream premature termination codon is indicated by the red **X**).

At the protein level, western blot analysis detected only full-length FPC in kidneys of normal mice. In contrast, no protein bands of any size were observed in immunoblots of *cyli/cyli* mouse kidney protein extracts. This observation is consistent with the reduced levels of *Pkhd1* gene expression demonstrated by qRT-PCR analysis of *cyli/cyli* kidney and leaves open the possibility that variant transcripts give rise to very low abundance isoforms of functional FPC, undetectable on western blots but capable of preventing a cystic kidney phenotype. Previous polysome-based analyses indicate that the *Pkhd1* splice variants are indeed translated in normal kidney [16].

While the basis for *PKHD1/Pkhd1* species-specific differences in renal disease expression is not fully understood, differences in phenotypic severity between orthologous human disease and mouse models are not uncommon. For example, the *mdx* mouse model of Duchenne muscular dystrophy (DMD) has a mild phenotype compared to human DMD patients [40]. Although *mdx* mice exhibit muscle histopathology, elevated plasma pyruvate kinase and creatine kinase levels and muscle weakness similar to DMD patients, *mdx* mice are viable and fertile [40] whereas human DMD is a fatal degenerative muscle disorder [41]. Mouse models of human cystic fibrosis (CF) deficient for cystic fibrosis transmembrane conductance regulator (*CFTR*) also fail to fully recapitulate the human disease [42]. More than 10 CF mouse models have been created and although they exhibit fluid secretion defects and develop severe intestinal and mild pancreatic disease, they fail to develop the signature lung infections that are the major cause of mortality in human CF [43-46]. The basis for this species-specific difference in lung phenotype involves pH differences in the airway surface liquid (ASL) in human CF patients *vs*. mouse models [47].

We speculate that the minimal renal cystic disease in mouse *Pkhd1* models reflects a combination of mechanisms. One mechanism, exemplified by the *cyli/cyli* mutant characterized in the present study, is *genetic* in nature and involves the *Pkhd1* mutation itself triggering NAS to drive increased proportional representation of *Pkhd1* mRNAs lacking mutated exon 48 sequences, which direct translation of low-abundance but functionally competent FPC isoforms, thus preventing kidney disease. A second mechanism, reflecting *molecular interactions*, which may be dictated by genetic background rather than *Pkhd1* genotype *per se*, might act to modulate the degree to which a kidney phenotype is expressed. Variations in molecular interactions could also account for the well-documented effects of genetic background on kidney phenotypes displayed by multiple mouse *Pkhd1* models [9, 48, 49]. Finally, we speculate that FPC functions differently in regulatory pathways in the mouse and human. Mice homozygous for the *Pkhd1* exon 67 deletion (*Pkhd1*^*del67*^), which removes most of the FPC carboxy terminus domain, have no renal or biliary phenotype [14], whereas mice lacking virtually the entire *Pkhd1* locus (exons 3 through 67) express the hepatobiliary lesion, but only a minor renal phenotype in older mice [17, 50]. The corresponding defects in human patients, involving loss of a functional carboxy terminus [39] or large deletions intragenic deletions [51, 52] are associated with both the hepatobiliary lesion and severe renal cystic disease.

As noted above for CF, identifying specific mechanisms underlying discordant phenotypes between human genetic disease and orthologous mouse mutant models can yield valuable insights into novel therapeutic targets and potential treatment strategies [47]. Although the physiological function of FPC remains undefined, we anticipate that continued study of *Pkhd1* mutant mouse models will increase understanding of the mechanism(s) underlying mouse resistance to the severe renal disease that characterizes human ARPKD. Defining such mechanisms, in turn, could yield potential new targets for prevention and treatment of this devastating disease.

## Acknowledgments

The authors thank members of the Guay-Woodford laboratory for helpful advice. Adam Richman (Center for Translational Research) assisted with writing and editing of the manuscript and Amber K. O’Connor (akoWriting LLC) provided editorial assistance. We thank Trenton R. Schoeb, DVM, PhD (UAB Comparative Pathology Laboratory) for assisting with the histopathological analysis.

The authors also thank members of the Eicher laboratory, including Linda L. Washburn for help throughout the project, Lisa Somes for maintaining the D.B/11Ei-*cyli* strain and providing mice for pathology studies, and Leona Gagnon and Andrew Rechnagle for isolating DNA and establishig DNA plates for mapping. In addition, we thank Douglas McMinimy, The Jackson Laboratroy, for conducting a genome scan to determine the chromosomal position of *cyli*.

